# Residual blinking-driven channel alignment for multicolor single-molecule localization microscopy

**DOI:** 10.64898/2025.12.23.696332

**Authors:** Fen Hu, Jianyu Yang, Mingxin Chen, Zhao Xie, Shuai Liu, Dan Ding, Leiting Pan, Jingjun Xu

## Abstract

Given the nanometer-scale resolution of single-molecule localization microscopy (SMLM), sequential multicolor SMLM is inherently challenged by inter-channel misalignment arising from unpredictable sample drift during channel switching. Traditional fiducial-based strategies are limited by added complexity in sample preparation. Here, we report a residual blinking-driven method for channel alignment in multicolor SMLM, termed ReBling alignment. By exploiting residual blinking events under dual-wavelength excitation, our method extracts shared localizations as intrinsic registration anchors and performs accurate cross-correlation-based alignment. We validated ReBling alignment across biological systems featuring distinct spatial patterns, including continuous patterned structures (mitochondria-microtubule co-imaging, dual-labeled microtubules), and discrete point-cluster assemblies (dual-color nuclear pore complex components), demonstrating its broad applicability and reliability. These results establish ReBling alignment as a practical, versatile, fiducial-free solution for accurate channel registration in sequential multicolor SMLM, enabling more precise and reliable characterization of complex cellular architecture and molecular interactions.

## Introduction

Single-molecule localization microscopy (SMLM) has emerged as a powerful technique for visualizing biological structures with nanometer-scale resolution ^1–4^. Particularly, multi-color SMLM, enables spatial mapping of molecular organization and interactions within cells ^5,6^. To achieve multi-color imaging, two primary strategies are typically employed: simultaneous and sequential acquisition ^7,8^. Simultaneous acquisition collects fluorescence signals from multiple color channels concurrently, utilizing approaches such as alternating activation lasers ^9^, optical splitting with dichroic mirrors and filters ^10,11^, spectral dispersion via prisms or gratings ^12–14^, spectral demixing algorithm ^15^ and point spread function (PSF) engineering ^16^. While this method allows for rapid imaging, it is often prone to significant crosstalk and interference between different channels, which may lead to artifacts or misinterpretation ^17^.

In contrast, sequential acquisition temporally separates the imaging of different channels, thereby minimizing spectral overlap and effectively avoiding crosstalk ^18–20^. However, it extends the total acquisition time and encounters challenges in correcting inevitable inter-channel drift ^7,8^. SMLM techniques such as direct stochastic optical reconstruction microscopy (dSTORM) rely on stochastic blinking of fluorophores, which repeatedly switch between fluorescent and non-fluorescent states ^18^. In sequential multicolor imaging, high-intensity illumination is typically applied for several minutes to quench fluorophores, ensuring a sparse blinking state appropriate for data acquisition ^21^. During this interval, thermal and mechanical instabilities can lead to unpredictable sample drift, which compromises accurate channel alignment and obscures the true spatial relationships among labeled targets.

Fiducial-based methods, typically employing multi-spectral fluorescent beads, offer a viable solution to this problem ^22,23^. While effective in principle, they complicate sample preparation and are therefore not routinely adopted in SMLM experiments. As a result, inter-channel alignment is commonly performed by directly applying the cumulative drift from the initial to the final segment measured in the first channel to the second. However, this approach neglects additional drift that may occur during the inter-channel waiting period, leading to systematic misalignments. In high-resolution SMLM (∼20 nm), this uncorrected drift can substantially impair the accuracy of nanoscale colocalization or interaction analyses.

To address these issues, we developed a fiducial-free registration method termed ReBling alignment that leverages residual blinking events induced under dual-wavelength excitation. By identifying shared localizations between channels and applying direct cross-correlation, ReBling alignment quantifies and corrects drift introduced during channel switching. This endogenous strategy bypasses the limitations of fiducial-based approaches and enables accurate alignment in sequential multicolor SMLM.

## Results

### Workflow of ReBling alignment for sequential two-color SMLM

The ReBling alignment method consists of two main parts: sequential data acquisition and computational post-processing for drift correction. During data acquisition, a standard dual-color sequential STORM SMLM protocol is employed. Excitation lasers at 647 nm and 561 nm are sequentially introduced into the sample through the back focal plane of a high numerical aperture objective, using a multicolor dichroic mirror that reflects both wavelengths (Figure 1a). Fluorescence emitted from the sample is collected by the same objective, transmitted back through the dichroic mirror, and filtered by a multicolor emission filter that rejects both excitation lasers (Figure 1a). The first imaging channel (Channel 1) is acquired under 647 nm excitation (Figure 1b), followed by the second channel (Channel 2) under 561 nm excitation (Figure 1c). Upon completion of the dual-color sequential imaging, a third acquisition step is performed in which both 647 nm and 561 nm excitation lasers are simultaneously applied. This generates a composite dataset (Channel 3) capturing “residual blinking signals” from both dye species without spectral discrimination, which arise from a small fraction of fluorophores that remain in reversible dark states or are not fully photobleached after the sequential acquisitions (Figure 1d). This third channel contains spatially co-localized signals from both primary channels and thus can serve as an endogenous reference for image registration.

**Figure 1.**
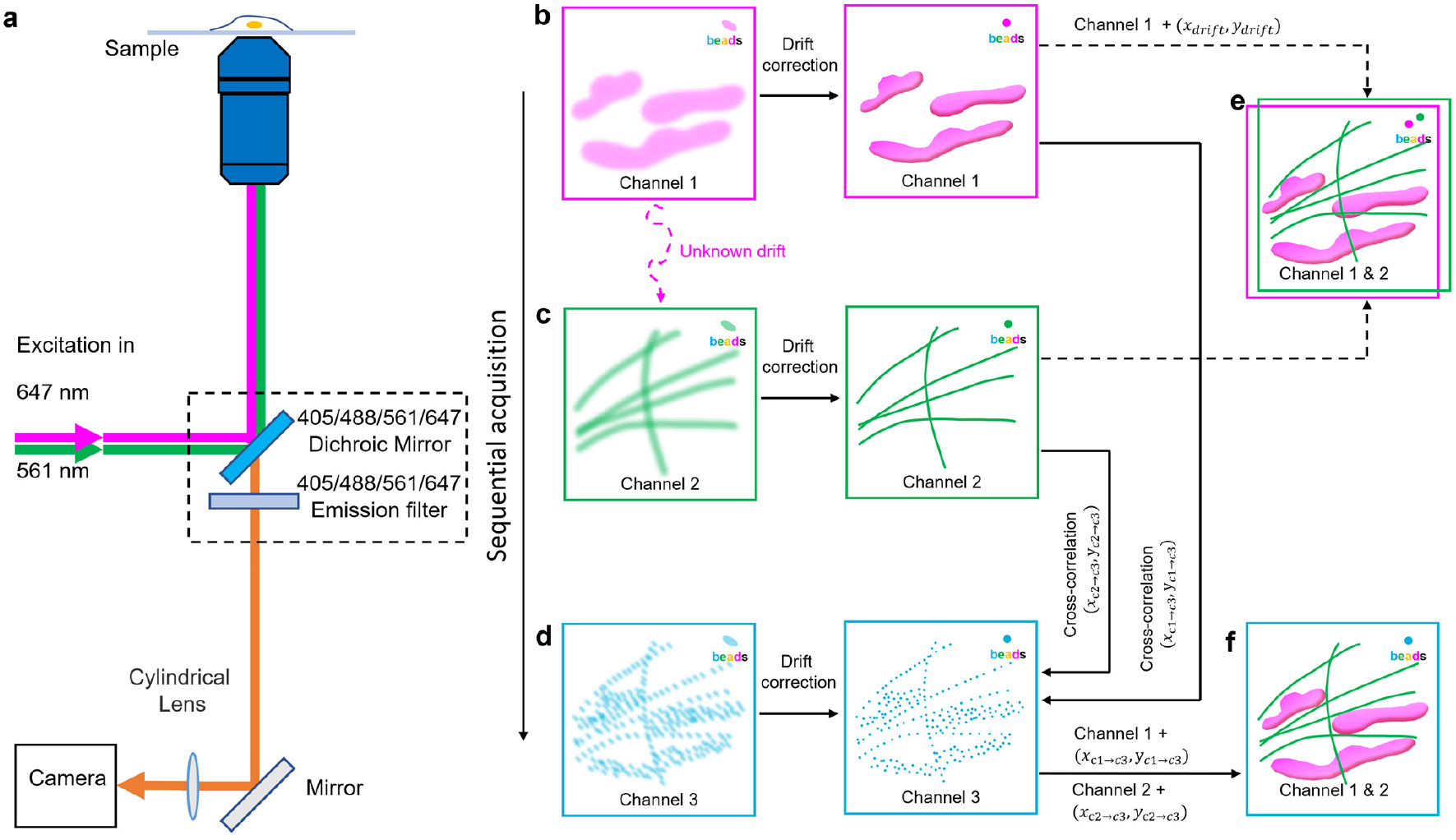
Workflow of ReBling alignment for inter-channel registration in sequential two-color SMLM. (a) Schematic of the dual-color STORM imaging system. Excitation lasers are directed into the objective lens via a multicolor dichroic mirror. Emitted fluorescence passes through the same dichroic and a multicolor emission filter, then shaped by a cylindrical lens to modulate the point spread function (PSF) before being recorded by the camera. (b) STORM image for Channel 1 under 647 nm excitation. Left: raw image; right: corresponding intra-channel drift-corrected image. (c) STORM image for Channel 2 under 561 nm excitation. Left: raw image; right: corresponding intra-channel drift-corrected image. (d) STORM image for Channel 3 under simultaneous 647 nm and 561 nm excitation, capturing residual blinking events from both fluorophores. Left: raw image; right: corresponding intra-channel drift-corrected image. (e) Schematic of conventional inter-channel alignment, where the cumulative drift of Channel 1 between its first and final segment is directly transferred to Channel 2. (f) Schematic of ReBling alignment, illustrating independent direct cross-correlation between Channels 1 and 3 and between Channels 2 and 3 to obtain relative drift vectors for registration.

In the post-processing step, single-molecule localization coordinates from all three channels are extracted using standard SMLM image processing algorithms. Each channel is then subjected to intra-channel drift correction via nearest paired cloud (NP-Cloud) ^24^, a fast and robust SMLM drift-correction method we have newly developed (Figure 1b-d).

Conventional inter-channel alignment typically transfers the total drift measured in Channel 1 to Channel 2, neglecting the untracked drift that may occur during the channel-switching interval required for fluorophore quenching (Figure 1e). To address this, the ReBling alignment approach leverages residual blinking signals captured under dual-wavelength excitation in Channel 3 as a shared reference. Direct cross-correlation ^9^ is independently performed between the dataset of Channel 1 and Channel 3 to obtain drift vectors (Δ_*x*1→3_, Δ_*y*1→3_), and between Channel 2 and Channel 3 (Δ_*x*2→3_, Δ_*y*2→3_). Briefly, single-molecule localization points were binned into two-dimensional density maps for each channel. The cross-correlation function between the two maps was then computed across a range of spatial displacements, and the peak position of this function identifies the relative shift between channels. Since Channel 3 contains overlapping localizations from both primary channels, these relative transformations are used to register Channel 1 and Channel 2 by referencing their positions to a common coordinate framework (Figure 1f). This strategy provides a reliable solution for inter-channel alignment in sequential two-color SMLM, thereby facilitating downstream analysis of nanoscale spatial relationships.

### Performance of ReBling alignment on dual-color SMLM imaging of microtubules and mitochondria

To evaluate the performance of ReBling alignment, we applied it to dual-color SMLM imaging of microtubules and mitochondria in Cos-7 cells, two well-defined, “patterned” cellular structures. Accurate resolution of their spatial relationships is essential, as their interaction underlies intracellular energy distribution and metabolic regulation ^25,26^. To visualize them, α-tubulin was labeled with CF583R-conjugated secondary antibodies, and the mitochondrial outer membrane protein TOMM20 was labeled with Alexa Fluor 647-conjugated secondary antibodies, respectively. Besides, multi-spectral fiducial fluorescent beads were immobilized on the coverslip as an external registration benchmark for registration validation.

Sequential imaging was conducted by first acquiring ∼40,000 frames under 647 nm excitation (Channel 1; Figure 2a), followed by another ∼40,000 frames under 561 nm excitation (Channel 2; Figure 2b). ReBling alignment was then applied through a third acquisition of 30,000∼40,000 frames under simultaneous 647 nm and 561 nm excitation to capture residual blinking signals (Channel 3; Figure 2c).

**Figure 2.**
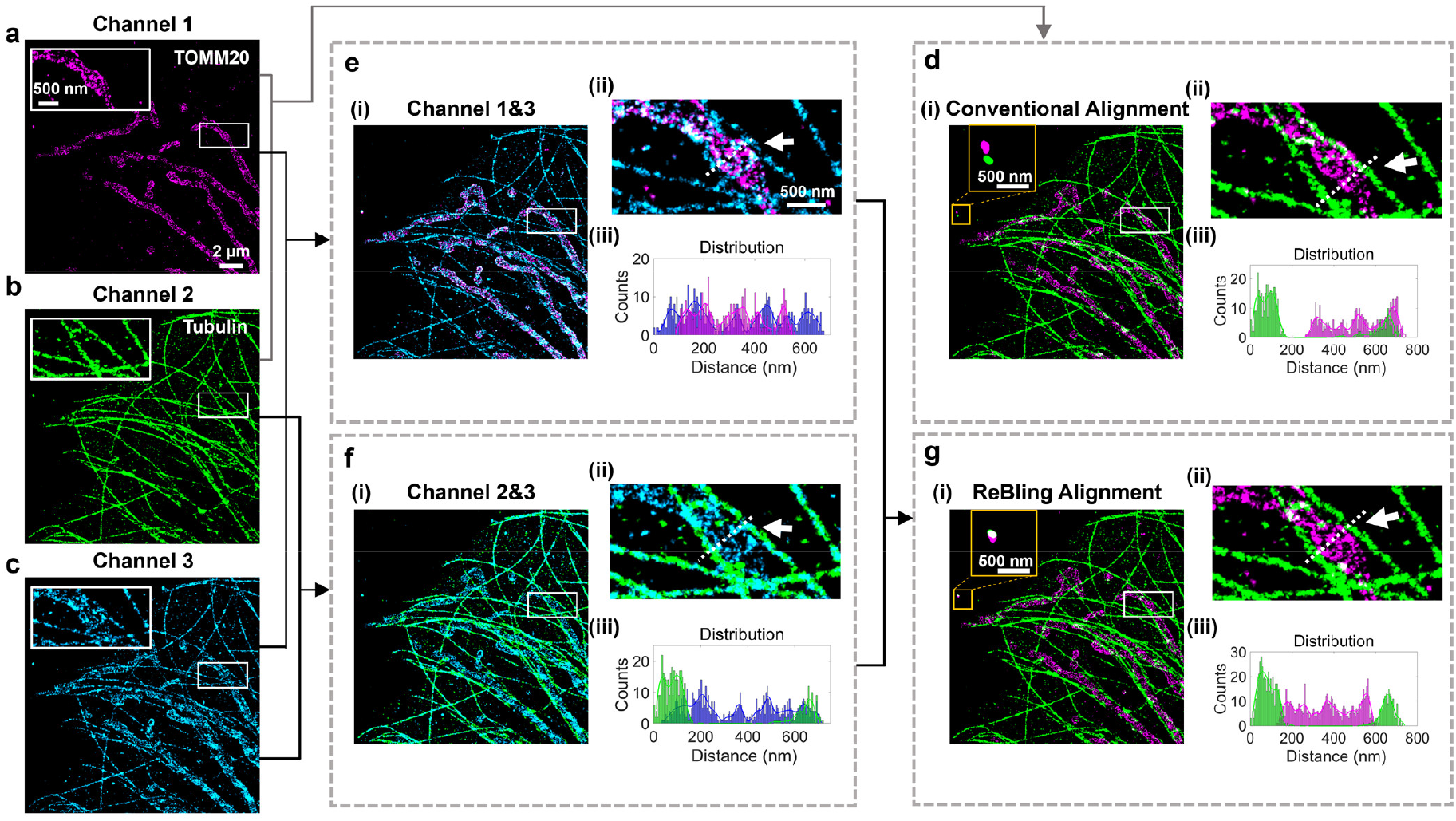
Alignment of dual-color STORM images of microtubules and mitochondria in Cos-7 cells. (a) STORM image of mitochondria in Channel 1, immunolabeled with Alexa647-conjugated secondary antibodies. (b) STORM image of microtubules in Channel 2, immunolabeled with CF583R-conjugated secondary antibodies. (c) STORM image of Channel 3, generated by simultaneous excitation at 647 nm and 561 nm, utilizing residual blinking. (d) Conventional alignment of Channels 1 and 2. The enlarged yellow box with fluorescent beads (i), the zoomed-in white box region (ii), and the localization count profiles along the white dashed line (iii), are shown. (e)Alignment of Channels 1 and 3 via direct cross-correlation (i). The zoomed-in region in the white box (ii) and corresponding localization count profiles (iii), are shown. (f) Alignment of Channels 2 and 3 presented similarly as in (e). (g) Alignment of Channels 1 and 2 using ReBling alignment. The enlarged view of the fluorescent beads in the yellow box (i), the zoomed-in white box (ii), and the localization count profiles (iii), are shown.

For conventional alignment, after intra-channel drift corrections by NP-Cloud, the cumulative drift of Channel 1 from the initial to the final segment was directly transferred to Channel 2 (Figure 2d). However, fiducial fluorescent beads exhibited a noticeable mismatch between the two channels (yellow box in Figure 2d(i)), indicating a substantial registration error. This misalignment led to an apparent overlap between a microtubule and a mitochondrion in the magnified local view (Figure 2d(ii), d(iii)), which may misrepresent their true spatial relationship.

For ReBling alignment, direct cross-correlation was performed between Channel 1/2 and 3, yielding relative displacement vectors used to register Channel 1 to 3 (Figure 2e), and Channel 2 to 3 (Figure 2f), respectively. The alignment between Channels 1 and 2 was then inferred through their respective registrations to Channel 3 (Figure 2g). This approach resulted in precise colocalization of fiducial signals (Figure 2g(i)) and a corrected spatial arrangement in which a mitochondrion was clearly resolved between two adjacent microtubules (Figure 2e(ii,iii), f(ii,iii), g(ii,iii)). These results underscore the efficacy of ReBling alignment in complex multi-component patterned structures where precise spatial relationships are critical for functional interpretation.

### Performance of ReBling alignment on dual-labeled SMLM imaging of microtubules

To validate the registration accuracy of ReBling alignment, we next performed a dual-labeled SMLM experiment in which α-tubulin was simultaneously labeled with Alexa Fluor 647- and CF583R-conjugated secondary antibodies in Cos-7 cells. As the same target is imaged in both channels, their merged signals are expected to show perfect spatial overlap.

Similarly, we compared the performance of conventional dual-channel alignment with that of ReBling alignment. In the conventional approach (Figure 3a,b,d), magnified views revealed pronounced misalignment between the two channels, as evidenced by shifted fiducial marker (yellow box in Figure 3d(i)), spatially offset microtubule filaments (Figure 3d(ii)), as well as discrepancies in localization count profiles (Figure 3d(iii)). These results indicate that conventional alignment fails to compensate for inter-channel drift accumulated during sequential acquisition.

**Figure 3.**
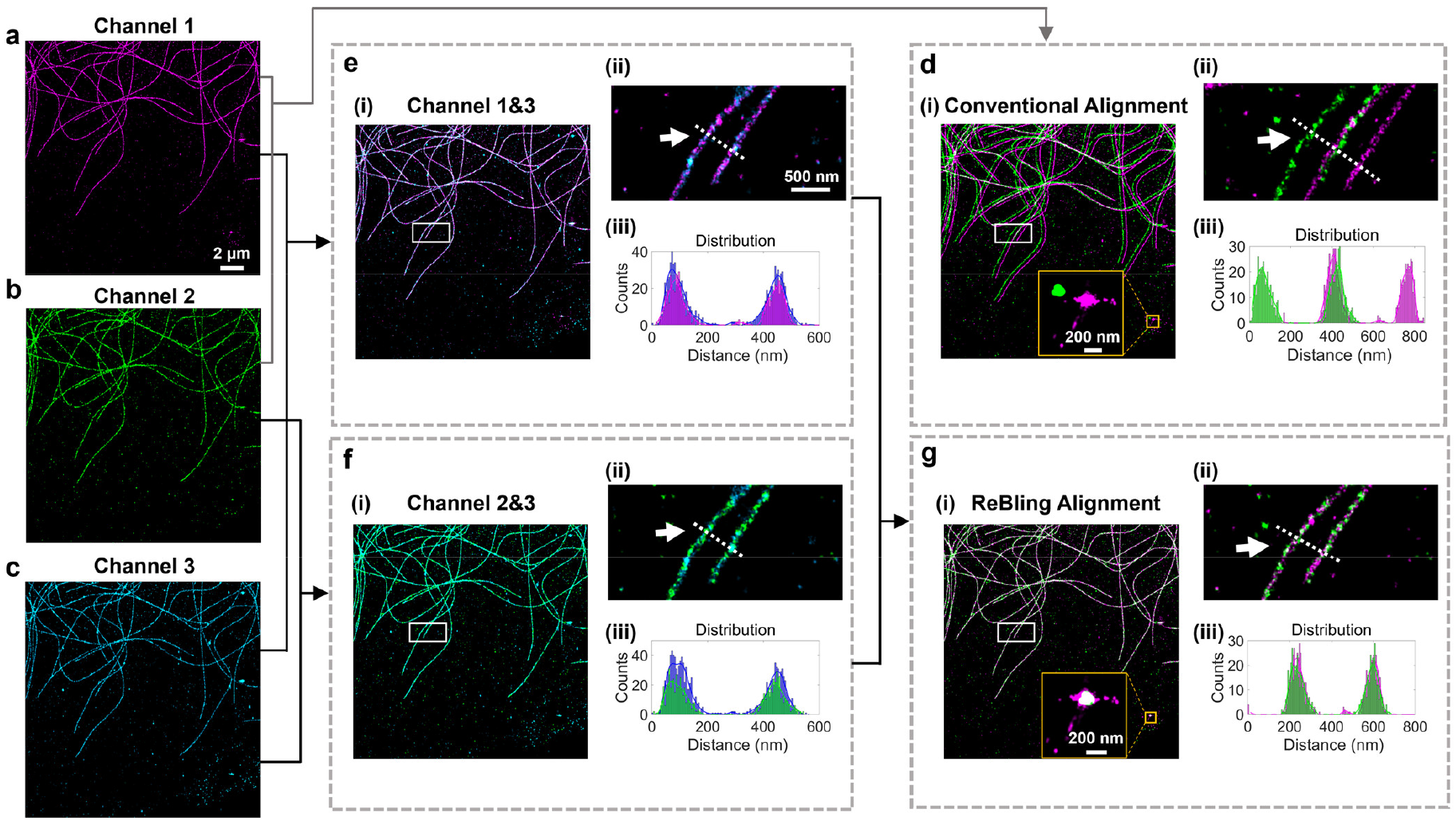
Alignment of two-color STORM images of microtubules in Cos-7 cells. (a) STORM image of microtubules in Channel 1, immunolabeled with Alexa647-conjugated secondary antibodies. (b) STORM image of microtubules in Channel 2, immunolabeled with CF583R-conjugated secondary antibodies. (c) STORM image of microtubules in Channel 3, generated by simultaneous excitation at 647 nm and 561 nm, utilizing residual blinking. (d) Conventional alignment of Channels 1 and 2. The yellow box highlights a fluorescent bead adhered to the substrate, with enlarged view shown alongside (i). The zoomed-in region of microtubules within the white box (ii), and the localization count profiles along the white dashed line (iii), are also shown. (e) Alignment of Channels 1 and 3 through direct cross-correlation (i). The zoomed-in region in the white box (ii), and corresponding localization count profiles along the white dashed line (iii), are shown. (f) Alignment of Channels 2 and 3, processed as in (e). (g) Alignment of Channels 1 and 2 using the ReBling alignment. Enlarged view of the fluorescent bead in the yellow box (i), zoomed-in white-box region (ii), and corresponding localization count profiles (iii), are shown.

By contrast, application of ReBling alignment (Figure 3c,e,f,g) yielded marked improved outcomes, as evidenced by co-localized fiducial signals in both channels (Figure 3g(i)), near-perfect spatial overlap of microtubule structures (Figure 3g(ii)), and coincident peaks of the localization count profiles (Figure 3g(iii)). These results further demonstrate that ReBling alignment effectively corrects inter-channel drift and achieves accurate registration in sequential dual-color SMLM.

### Performance of ReBling alignment on two-color SMLM imaging of nucleoporins Nup98 and Nup62

Finally, to assess the applicability of ReBling alignment to “point-cluster” SMLM data, we employed the method to resolve the spatial relationship between two critical components of the nuclear pore complex (NPC): Nup98 and Nup62, in U2OS cells. The NPC is a large macromolecular assembly composed of multiple nucleoporins (Nups) arranged in defined subcomplexes ^27^. Among them, Nup62 is a central channel nucleoporin located predominantly in the axial core of the pore, while Nup98 distributed toward the peripheral filaments or cytoplasmic rings ^28^.

In datasets processed using conventional alignment (Figure 4a,b,d), the spatial arrangement of Nup98 and Nup62 appeared deviated from established NPC structural models. Specifically, the zoomed-in view of a representative NPC showed a clear spatial separation between the two nucleoporins (yellow box in Figure 4d(i,ii)), indicative of poor colocalization. This observation was further supported by two-dimensional pairwise cross-correlation analysis ^29,30^, which exhibited a cross-correlation peak value away from zero distance (Figure 4d(iv)), suggesting a misalignment between the channels.

**Figure 4.**
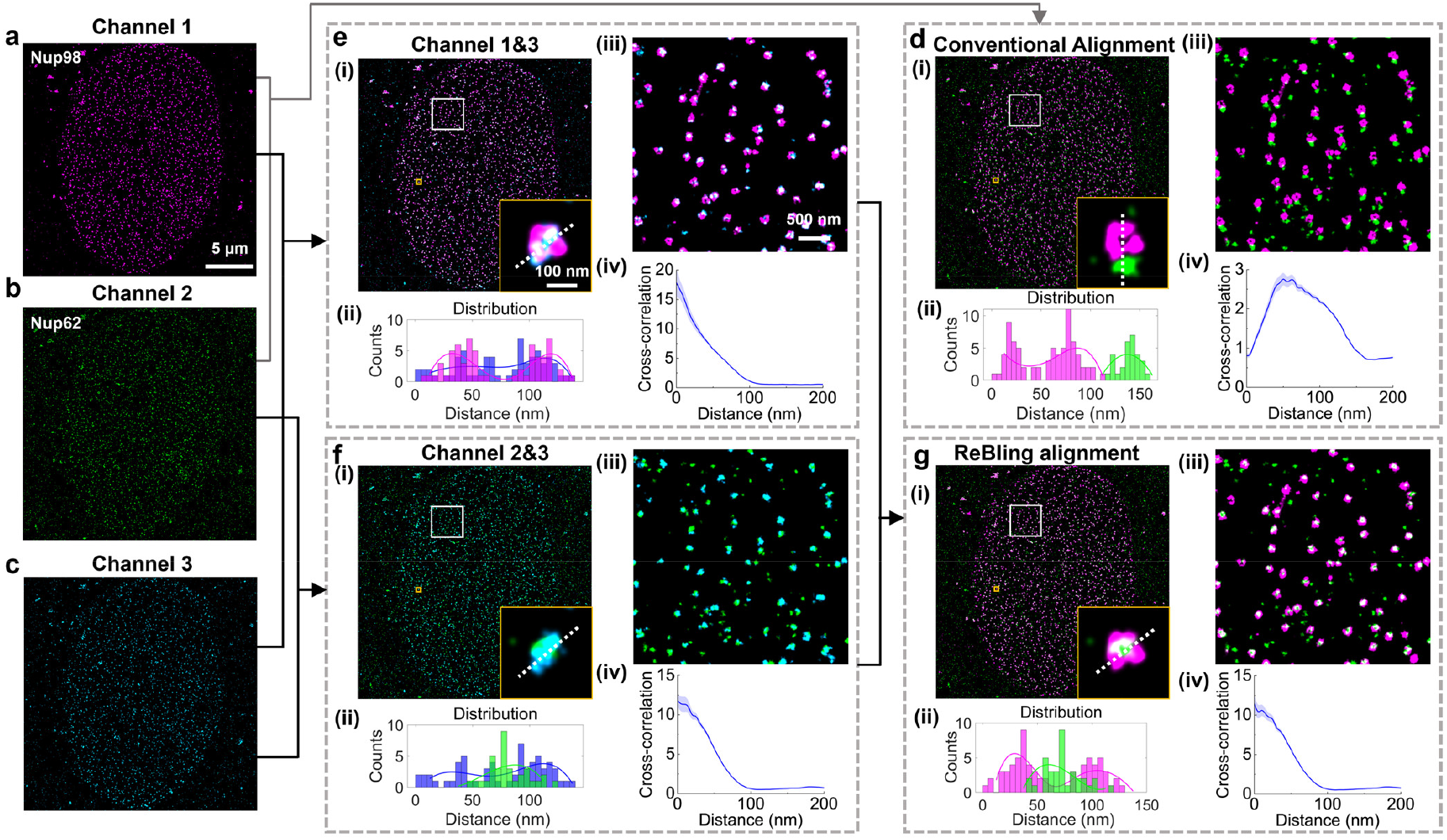
Alignment of two-color STORM images of Nup98 and Nup62 in U2OS cells. (a) STORM image of Nup98 in Channel 1, immunolabeled with Alexa647-conjugated secondary antibodies. (b) STORM image of Nup62 in Channel 2, immunolabeled with CF583R-conjugated secondary antibodies. (c) STORM image of Channel 3, generated by simultaneous excitation at 647 nm and 561 nm, utilizing residual blinking. (d) Conventional alignment of Channels 1 and 2. The enlarged yellow box highlights an individual NPC (i), and the localization count profiles along the white dashed line (ii). The enlarged view in the white box (iii) and colocalization assessment by cross-correlation analysis (iv), are also shown. (e) Alignment of Channels 1 and 3 via direct cross-correlation. The enlarged views within the yellow box along with the localization count profiles (ii), are shown. The enlarged views within the white box (iii) along with colocalization assessment result (iv), are also shown. (f) Alignment of Channels 2 and 3 performed similarly as in (e). (g) ReBling alignment of Channels 1 and 2. The enlarged yellow box highlights an individual NPC (i), and localization count profiles (ii), are shown. The enlarged view in the white box (iii) and colocalization assessment result (iv), are also shown.

In contrast, application of ReBling alignment (Figure 4c,e,f,g) recovered the expected spatial arrangement, with Nup62 clearly localized in the central region of the NPC, encircled by Nup98 signals (Figure 4g(i,ii)), in line with known NPC architecture. Cross-correlation analysis confirmed this spatial correspondence, yielding a cross-correlation peak value exceeding 1 at zero distance (Figure 4g(iv)). These results showcase the superior performance of ReBling alignment in restoring nanoscale accuracy in structures with “point-cluster” distributions.

## Discussion

In this study, we developed ReBling alignment, a novel registration strategy designed to overcome the critical bottleneck of inter-channel alignment in sequential dual-color SMLM. Traditional fiducial marker-based registration methods ^22,23^, though generally reliable, require the introduction of external markers such as multi-color fluorescent beads into the sample. Furthermore, controlling the spatial distribution of fiducials is difficult: sparse markers may be absent from the field of view, while dense markers can interfere with biological structures of interest, causing imaging artifacts thus limiting their practical application in SMLM. Although alternative approaches such as rapid channel switching to minimize delay, or pre-quenching fluorescence of the second channel have been employed, inter-channel drift remains a persistent challenge. This issue becomes increasingly crucial as higher resolution impose stricter demands on drift correction to ensure accurate analysis of molecular organization and interactions.

Our proposed method, ReBling alignment, offers a conceptually different solution by exploiting residual blinking events under simultaneous dual-wavelength excitation to generate shared localizations as intrinsic registration anchors (Figure 1). By performing direct cross-correlation on these co-occurring localizations, the method accurately determines and corrects the untracked drift during channel switching. This marker-free approach circumvents the drawbacks of fiducial-based methods, making it especially advantageous for densely labeled or structurally delicate specimens. Importantly, ReBling alignment is fully compatible with standard STORM protocols and requires no hardware modifications, allowing for seamless integration into existing imaging workflows.

SMLM data exhibit diverse spatial organizations depending on the biological targets, ranging from discrete point-cluster distributions represented by membrane proteins to continuous patterned structures such as cytoskeletal filaments or organelles. These varied spatial modes necessitate tailored analysis approaches, including clustering, morphological reconstruction, and colocalization analysis ^31–33^. The utility and effectiveness of ReBling alignment were validated across multiple cellular targets encompassing these varied spatial organizations. In dual-color SMLM imaging of microtubules and mitochondria (patterned structures), the method consistently achieved precise channel registration (Figure 2 and 3). It was further employed to visualize the colocalization of nuclear pore complex proteins Nup98 and Nup62 (point-cluster assemblies) (Figure 4). These results highlight the broad applicability of ReBling alignment to complex, multi-component subcellular structures.

## Conclusion

In summary, ReBling alignment provides a simple yet powerful solution for inter-channel alignment in sequential dual-color SMLM super-resolution microscopy. Its marker-free nature, ease of implementation, and resolution-matched accuracy make it a practical tool for super-resolution imaging. As SMLM continues to advance toward increasingly precise analysis of molecular interactions and spatial organization, ReBling alignment offers timely and enabling support for the reliable interpretation of nanoscale biological systems.

## Materials and Methods

### Reagents

EM-grade paraformaldehyde and glutaraldehyde were purchased from Electron Microscopy Sciences (USA). Mouse monoclonal antibodies against α-tubulin, rabbit monoclonal antibodies against TOMM20 were sourced from Nano-Microimaging Biotechnology (China). Mouse monoclonal antibodies of Nup98 were from Cell Signaling Technology (USA). Rabbit monoclonal antibodies of Nup62 were from Becton, Dickinson and Company (USA). Alexa Fluor 647 AffiniPure donkey anti-mouse/rabbit IgG (H+L) were from Invitrogen (USA). CF583R AffiniPure donkey anti-mouse IgG (H+L) were from Biotium (USA). Bovine serum albumin (BSA), Triton X-100, and other general reagents were obtained from Sigma-Aldrich (USA).

### Cell Culture

Cos-7 or U2OS cells were cultured in Dulbecco’s Modified Eagle Medium (DMEM) supplemented with 10% fetal bovine serum (FBS) in a humidified CO_2_ incubator with 5% CO_2_ at 37 °C, following standard tissue-culture protocols.

### Sample Preparation

Acid-washed 12 mm glass coverslips were washed with deionized water and dried before being placed in 24-well plates. For imaging of target proteins, cell samples were fixed with 3% w/v paraformaldehyde and 0.1% w/v glutaraldehyde in PBS for 20 min. After reduction with a freshly prepared 0.1% sodium borohydride solution in PBS, the samples were permeabilized and blocked in a blocking buffer (3% w/v BSA, 0.5% v/v Triton X-100 in PBS) for 20 min. Afterward, the cells were incubated with the primary antibody described above in blocking buffer for 1 h. After washing in washing buffer (0.2% w/v BSA and 0.1% v/v Triton X-100 in PBS) for three times, the cells were incubated with the secondary antibody for 1 h at room temperature. Then, the samples were washed three times with washing buffer before mounted for imaging.

### STORM super-resolution microscopy

For STORM imaging, samples on 12-mm coverslips were mounted on freshly cleaned 22 mm × 60 mm rectangular glass slides, following established protocols ^34,35^. Imaging was performed on a commercial STORM system (STORM Ultra300, Nano-Microimaging Biotechnology, China), which is based on an inverted optical microscope (Ti-E, Nikon, Japan) equipped with an EMCCD (iXon Ultra 897, Andor, UK) and a 100× oil-immersion objective (CFI Plan Apochromat λ, numerical aperture = 1.49, Nikon, Japan). Excitation was provided by 647-nm and 561-nm lasers sequentially coupled into the back focal plane of the objective. A multicolor dichroic mirror reflecting both wavelengths was used for excitation, and fluorescence signals were collected through the same objective and passed through a multiband emission filter that blocked the excitation lines (Figure 1a). High-power excitation (∼2 kW cm^-2^) drove most fluorophores into dark states while sparsely activating single emitters, and a low-intensity 405-nm laser (typically 0-1 W cm^-2^) was applied concurrently to stochastically reactivate fluorophores, ensuring that only a small, optically resolvable fraction of fluorophores were in the emitting state at any given instant. Images were recorded at 100 frames per second, with ∼40,000 frames typically recorded per dataset. Single-molecule localizations were determined using a previously described algorithm ^34,35^. Lateral drift during acquisition was corrected by NP-Cloud ^24^, providing a stable basis for subsequent channel registration. For ReBling alignment, relative displacement vectors between Channels 1/2 and the reference Channel 3 were obtained by cross-correlation ^9^, enabling indirect registration of Channels 1 and 2 through their independent alignments to Channel 3 (Figure 2).

### Two-dimensional pairwise cross-correlation analysis

Two-dimensional pairwise cross-correlation analysis ^29,30^ was conducted by calculating the pairwise intermolecular distances between single molecules identified in the two channels. To assess the significance of spatial correlation, the resulting distance distribution was normalized against those derived from multiple sets of randomly distributed molecules within the same region. The normalized cross-correlation amplitude at given displacements reflects the degree of spatial correlation between the channels, with values greater than 1 indicating positive correlation and values less than 1 indicating anti-correlation.

## Authors’ Contributions

L.P. conceived the research. L.P., F.H. and J.Y. designed the experiments. J.Y. and M.C. performed the experiments. M.C., Z.X., and S.L. analyzed the data. D.D. contributed to data validation and discussion. F.H. and L.P. wrote the manuscript. L.P. and J.X. supervised the work.

## Availability of data and materials

The data that support the findings of this study are available from the corresponding author upon reasonable request.

